# Optimization of On-Bead Emulsion Polymerase Chain Reaction Based on Single Particle Analysis

**DOI:** 10.1101/2020.07.10.159483

**Authors:** Ryan H.P. Siu, Yang A. Liu, Kaitlin H. Y. Chan, Clara Ridzewski, Liane Siu Slaughter, Angela R. Wu

**Author notes:** These authors contributed equally (Ryan H.P. Siu, Yang A. Liu, Kaitlin H. Y. Chan).

## Abstract

Emulsion polymerase chain reaction (ePCR) enables parallel amplification of millions of different DNA molecules while avoiding bias and chimeric byproducts, essential criteria for applications including next generation sequencing, aptamer selection, and protein-DNA interaction studies. Despite these advantages, ePCR remains underused due to the lack of optimal starting conditions, straightforward methods to evaluate success, and guidelines for tuning the reaction. This knowledge has been elusive for bulk emulsion generation methods, such as stirring, the only methods that can emulsify libraries of ≥10^8^ sequences within minutes, because these emulsions have not been characterized in ways that preserve the heterogeneity that defines successful ePCR. Our study systematically quantifies the influence of tuning individual ePCR parameters by single particle analysis, which preserves this heterogeneity. We combine ePCR with magnetic microbeads and quantify the amplification yield via qPCR and the proportion of clonal and saturated beads via flow cytometry. Our single particle level analysis of thousands of beads resolves two key criteria that define the success of ePCR: 1) whether the target fraction of 20% clonal beads predicted by the Poisson distribution is achieved, and 2) whether those beads are partially or maximally covered by amplified DNA. We found that increasing the polymerase concentration 20-fold in ePCR over recommended concentrations for conventional PCR was necessary to obtain sufficient PCR products. Surprisingly, dramatical increases in the concentrations of reverse primer and nucleotides recommended in literature gave no measurable change in outcome. We thus provide evidence-based starting conditions for effective and economical ePCR for real DNA libraries and a straightforward workflow for evaluating the success of tuning ePCR prior to downstream applications.

---

**Scheme 1.**
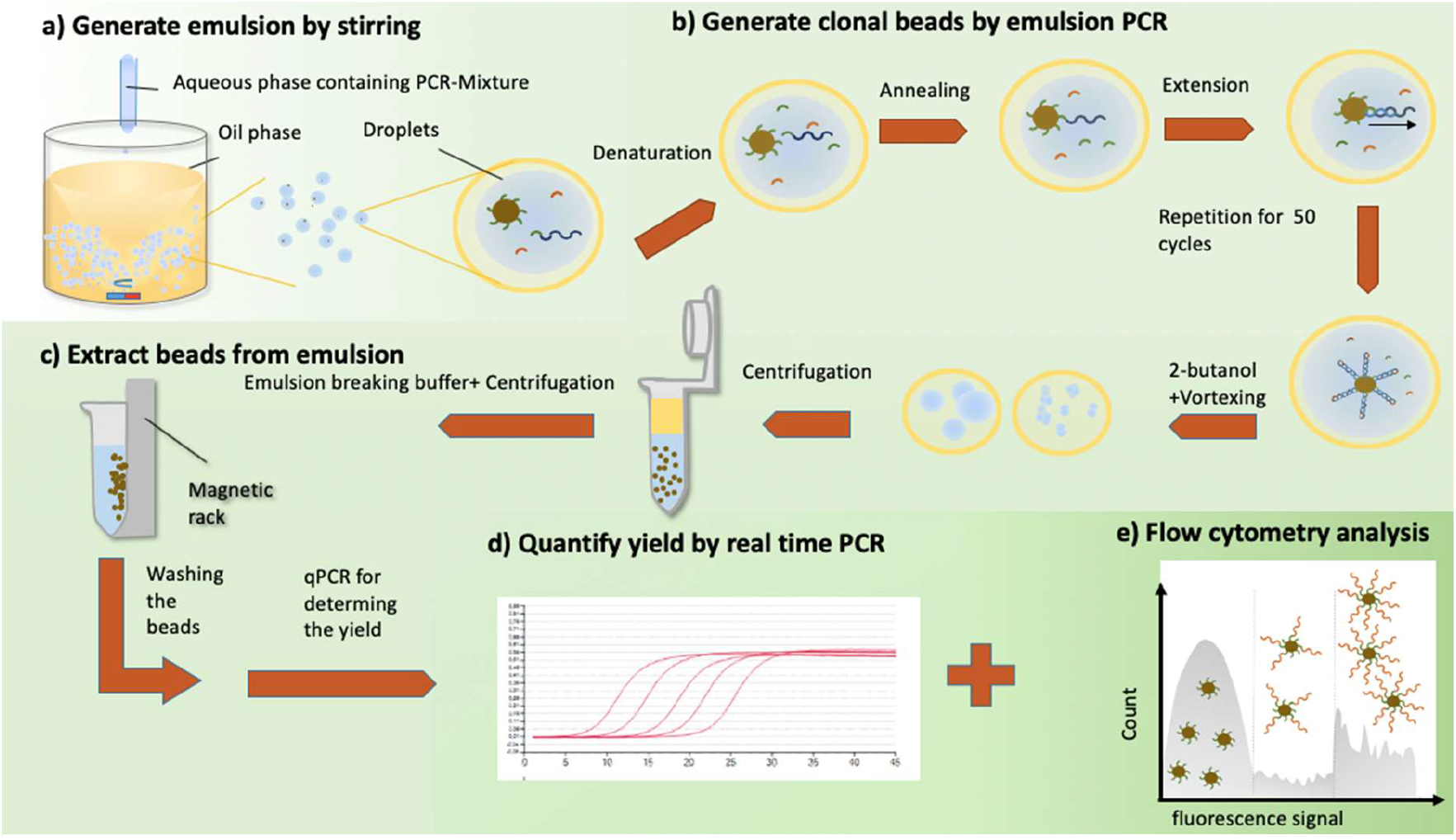
Overview of the workflow for optimizing on-bead emulsion PCR. a) Generation of emulsion by stirring. b) Emulsion PCR gives blank, clonal, and saturated beads. c) Beads are extracted and purified from the emulsion by centrifugation followed by magnetic separation. d) Quantitative PCR assay. We subject the extracted beads to qPCR to determine the yield of the ePCR and enrichment over input. e) Flow cytometry analysis. We pass the beads purified from emulsions to the flow cytometer and check the percentage of clonal beads and the relative number of amplicons per bead.

## 1. Introduction

Emulsion polymerase chain reaction (ePCR) is a version of PCR where an aqueous phase PCR mixture and an oil phase supplemented with surfactants are emulsified to form water-in-oil drops. Thousands to millions of DNA templates are amplified inside these drops in isolated but parallel reactions, as each drop serves as an isolated microreactor. Successful emulsion PCR gives numerous advantages over conventional, bulk PCR. First, separating template molecules avoids amplification bias.[1,2] Second, it reduces formation of unwanted, chimeric products caused by templates annealing to each other.[3] These features are required in applications where the relative abundance of different template molecules must be preserved, including whole genome sequencing, variant profiling, RNAseq, and aptamer selection.[4–9] For small scale applications with ≤40 μl of aqueous phase PCR such as droplet digital PCR (ddPCR), microfluidic devices are excellent for generating a user-desired number of highly uniform drops from small sample volumes,[10–13] necessary for accurate counting of target molecules.[10,14] However, microfluidics do not simultaneously offer the throughput and accessibility for ePCR of libraries comprising >10^8^ sequences, pertinent to the size of libraries in aptamer selections and next generation sequencing. In contrast, bulk methods for generating emulsions, such as stirring or vortexing, can generate millions to trillions of drops within minutes using common laboratory equipment.[14–17]

Despite its attractive advantages, ePCR remains underused for library amplification due to a lack of well-informed starting conditions, methods for evaluating success, and guidelines for optimizing the emulsion, especially for emulsions generated by bulk methods such as stirring and vortexing. Ideal ePCR requires heterogeneous distributions of reactants and templates in the drops such that 1) templates are segregated and each drop contains no more than one template, 2) every template-containing drop contains sufficient polymerase, primers, and nucleotides so that products can be amplified to completion. However, previous studies using end-point, ensemble-level analyses of the purified PCR products failed to elucidate these characteristics.[18–21] Thus, optimization of ePCR would be difficult According to Poisson statistics, there is a ‘sweet spot’ where the number of single-template drops is maximized with minimal number of multi-template droplets. This condition results in only ~20% of emulsion drops containing a template molecule. This ideal situation is heterogeneous; and only an analysis method of ePCR products that reflects this heterogeneity can determine the success of ePCR by which sufficient amplification and optimal template distribution have been achieved.

We present here an empirical strategy for evaluating the success of ePCR and a systematic study of the influence of tuning ePCR parameters while preserving the heterogeneous distributions of PCR components in the emulsion. In doing so, we define viable starting parameters that users can immediately apply for amplification of DNA libraries. We employed a strategy that reveals both the yield and distribution of PCR components in the drops by combining ePCR with magnetic microbeads. Forward primers are affixed to the surface of the magnetic microbeads to make forward-primer beads (FP-beads) so that all amplicons are immobilized onto the beads during ePCR. After clean-up, thousands of single beads per sample are analyzed by fluorescence flow cytometry to reveal the proportion of beads with zero, intermediate, and saturated numbers of amplicons. This heterogeneous distribution elucidates the relative rate at which beads co-occurred with template molecules and sufficient PCR reagents in the same drop, information that is inaccessible by ensemble characterization but necessary for tuning the protocol effectively. Using this strategy, we tuned the concentrations of polymerase, reverse primer, and deoxyribonucleotide triphosphates (dNTPs). The guidelines presented herein enable researchers to take advantage of the separation of templates in ePCR, and the scalability of generating emulsions by bulk methods. Our reaction and characterization strategy is easy to scale between thousands and hundreds of millions of parallel reactions. We provide a straightforward, versatile protocol for newcomers to ePCR. We also provide a robust flow cytometry-assisted single particle analysis methodology for ePCR researchers to evaluate their PCR product and optimize ePCR formulation according to their needs.

## 2. Materials and Methods

### 2.1 Materials and reagents

All DNA templates and primers were purchased from Integrated DNA Technologies. The oligonucleotide sequences are presented in the Supporting Information (Table S1). Dynabeads MyOne^™^ Carboxylic Acid and 100mM of each dNTP were purchased from Thermo Fisher Scientific. 1-Ethyl-3-(3-dimethylaminopropyl) carbodiimide (EDC) was purchased from Life Technologies/Thermo Fisher Scientific. Amino-PEG12 and 2-(N-morpholino) ethanesulfonic acid (MES) buffer (100 mM, PH 4.7) were purchased from Pierce Biotechnology. Amino-PEG12 was resuspended to 20 mM in DMSO before the first use. Span^®^ 80, Tween^®^ 80, Triton^™^ X100, N-hydroxysuccinimide (NHS) and light mineral oil were purchased from Sigma-Aldrich. GoTaq G2 Hot-start polymerase, 5X PCR Flexi Buffer and 2X GoTaq PCR Master Mix were purchased from Promega. An aliquot of ABIL^®^ EM 90 by Evonik was gifted to us from Professor Shuhuai Yao’s lab (see Acknowledgements).

### 2.2 Preparation of forward primer beads (FP-beads)

A previously published protocol was used to conjugate forward primers (FPs) onto the beads, enabling amplification of DNA attached to the beads.[15,22] Briefly, 5’-amino-modified FPs were covalently bonded to 1-μm MyOne carboxylic acid magnetic beads by coupling with N-hydroxysuccinimide (NHS) and 1-Ethyl-3-(3-dimethylaminopropyl)carbodiimide (EDC). The modified FP contained a polyethylene glycol (PEG_18_) spacer region between the amine group and the FP DNA sequence. Amino-modified PEG_18_ molecules were also coupled to the beads to separate neighboring FPs on the beads. The forward primer conjugated magnetic beads are referred as FP-bead.

### 2.3 Emulsion preparation (800 μl total emulsion volume)

Two oil phases were tested: ‘EM90-based oil’ with 3% (v/v) ABIL^®^ EM 90and 1% (v/v) Triton X-100 in mineral oil; ‘Span80-based oil’ with 4.5% (v/v) Span^®^ 80, 0.4% (v/v) Tween^®^ 80 and 0.05% (v/v) Triton^™^ X-100 in mineral oil. The aqueous phase consisted of 1X PCR Flexi Buffer, 2 pM single-stranded DNA template, ~3 x 10^5^/μl FP-beads, 25 mM MgCl_2_, 0.4 mM or 3.5 mM dNTP each, 2 μM or 10 μM GD1R and 0.025 U/μl or 0.5 U/μl GoTaq G2 Hot Start polymerase. We recommend sonicating the FP-beads prior to adding them to the aqueous phase, and to vortex the aqueous mixture before generating the emulsions.

To generate emulsions, 700 μl of oil phase was first added into a 0.5 dram shell vial (Kimble) with a PTFE 8 x 1.5mm magnetic stir bar (Cowie), and stirred at speed 3 on the Hotplate stirrer UC152 (Stuart) for 5 minutes. We chose this stirring speed to generate an ideal average drop size as calculated by the Poisson distribution (Figure S3, Supporting Information). Subsequently, 100 μl of aqueous phase was added dropwise over 30 seconds to the oil and the emulsion was stirred for at least another 5 minutes. Then the emulsion was distributed in 100μl aliquots into 8 PCR tubes. We then performed emulsion PCR under the following cycling conditions: 95 °C for 7 min, followed by 50 cycles of 95 °C for 30 sec, 65 °C for 30 sec and 72 °C for 75 sec and a post-cycling extension at 72 °C for 5 min. [18] A detailed protocol on how the beads are cleaned post-ePCR is in the supporting information.

### 2.4 Determining the yield of emulsion PCR by qPCR

Beads were evaluated at 1000 bead/well occupancy using the Kubo Quantagene q225 real-time PCR system (China) to determine the quantity of the amplified DNA sequences on beads. Since the DNA amplified during ePCR are attached to the beads, the total number of ssDNA quantified by qPCR must be scaled by the relative bead recovery after emulsion extraction (calculated as the ratio between the number of extracted beads and the number of beads added in the aqueous phase) to compensate for beads lost during extraction. The yield of emulsion PCR is reported as an amplification factor, calculated as follows:

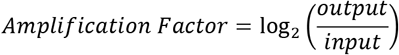

Output: Copy number of DNA product

Input: ssDNA input into the aqueous phase

### 2.5 Labelling bPCR and ePCR products with fluorescent probes

First, the PCR products were dehybridized by separating the beads from the supernatant, resuspending them in 0.1M NaOH, incubating them at 50°C for 5 minutes, and removing the supernatant. After repeating the suspension in NaOH, incubation, and separation once, the beads were washed three times with TE buffer. We then incubated the beads with 0.2 μM 5’ TYE^™^665–GD1R, the reverse primer-dye conjugate, at 60°C for 5 minutes. The beads were washed with PBSMCT buffer (DPBS with 2.5 mM MgCl_2_, 1 mM CaCl_2_, 0.05% Tween 20) before flow cytometry analysis.

### 2.6 Analysis of PCR products by flow cytometry

At least 10,000 beads per sample were analyzed by flow cytometry on a BD FACS Aria III (BD Biosciences). We used the 70 micron nozzle and no neutral density filter during analysis. The absorption, emission and bandpass wavelengths for the fluorescence channel were 650, 660, 660/20 nm respectively. The voltage settings for the photomultiplier tube (PMT) detectors were set to 300, 360 and 600V for the forward scattering, side scattering, and fluorescence, respectively, but small adjustments were made according to day-to-day variations of the instrument.

## 3. Results and Discussion

### 3.1 Study design and rationale

We first synthesized forward-primer beads (FP-beads) by covalently attaching forward primers to magnetic beads through amide bond-formation, an approach already established for on-bead ePCR. After synthesizing FP-beads, we first optimized non-emulsion on-bead PCR (bPCR) to achieve saturation of amplicons on beads. We performed bPCR with various cycle numbers to identify when a sufficient number of amplicons per bead is reached without non-specific PCR products (Figures S1 and S2.) Doing so defines the reference against which to judge whether on-bead ePCR products have reached completion.

We formed emulsions by stirring, which is both time-efficient and feasible by most laboratories without specialized equipment. We investigated two formulas, EM90-based and Span-80-based, that are common in the field and empirically determined their thermal stability from macroscopic and microscopic observations. (Figures S3 and S4, Supporting Information). Since we found that the EM90-based oil phase formula forms more heat-stable emulsions than Span80-based oil, all subsequent tests in this study use EM90-based oil.

We then tested different concentrations of polymerase, reverse primer, and nucleotides, starting with the concentrations recommended for regular PCR by the manufacturers of the polymerase. We tested the effect on yield and percentage of saturated beads by raising each component’s concentration as stated by another group of researchers who successfully developed an on-bead ePCR protocol for aptamer selection.[19] Since prior ePCR users reported using concentrations that are much higher than conventional PCR, we tested both higher and conventional concentrations.[23–26] Most ePCR applications increase the concentrations of each of these components by 4- to 20-fold compared with conventional PCR without documenting the change in outcome. We document the consequences of these extremes herein. Table 1 summarizes the different conditions that were tested in this study. Among the conditions we tested, we found that the higher concentration of polymerase (0.5 U/μl) produces 20.1% ± 4.0% clonal beads, an optimal proportion to generate monoclonal beads according to Poisson statistics. Surprisingly, using higher concentrations of reverse primer (10 μM) and nucleotides (3.5 mM) resulted in no significant improvements in both the proportion of monoclonal beads and the overall yield of ePCR products, suggesting that the higher concentrations used in prior studies expend reagents needlessly.

**Table 1.** Different Biochemical Parameters Tested in ePCR

| Condition | Polymerase | Reverse Primer | dNTP |
| --- | --- | --- | --- |
| **A** | 0.025 U/μl (Low) | 2 μM (Low) | 0.4 mM each (Low) |
| **B** | 0.5 U/μl (High) | 2 μM (Low) | 0.4 mM each (Low) |
| **C** | 0.5 U/μl (High) | 10 μM (High) | 0.4 mM each (Low) |
| **D** | 0.5 U/μl (High) | 2 μM (Low) | 3.5 mM each (High) |

### 3.2 Strategy for determining the degree of amplicon coverage on bead by flow cytometry

We defined the fluorescent signals of beads with no amplification products using FP-beads hybridized with unlabeled DNA sequences complementary to the forward primer. Beads with signal beyond this threshold were defined as clonal, meaning that they have amplification products on their surface, and gated them as clonal beads as shown in Figure 1A. To obtain signals defining saturated beads, which hold highest amount of amplicons achievable empirically, we performed bPCR using an optimized protocol to generate positive control saturated beads. We set the threshold to include the 90% of the positive control beads with the highest signal as shown in Figure 1B. We thus gated the ePCR beads into negative, clonal, and saturated beads as shown in Figure 1C. The test ePCR samples were analyzed by flow cytometry and characterized as shown in Figure 2. The flow cytometry gates are set according to control samples on the same day as measurements of ePCR samples to account for day-to-day variations of the instrument.

**Figure 1.**
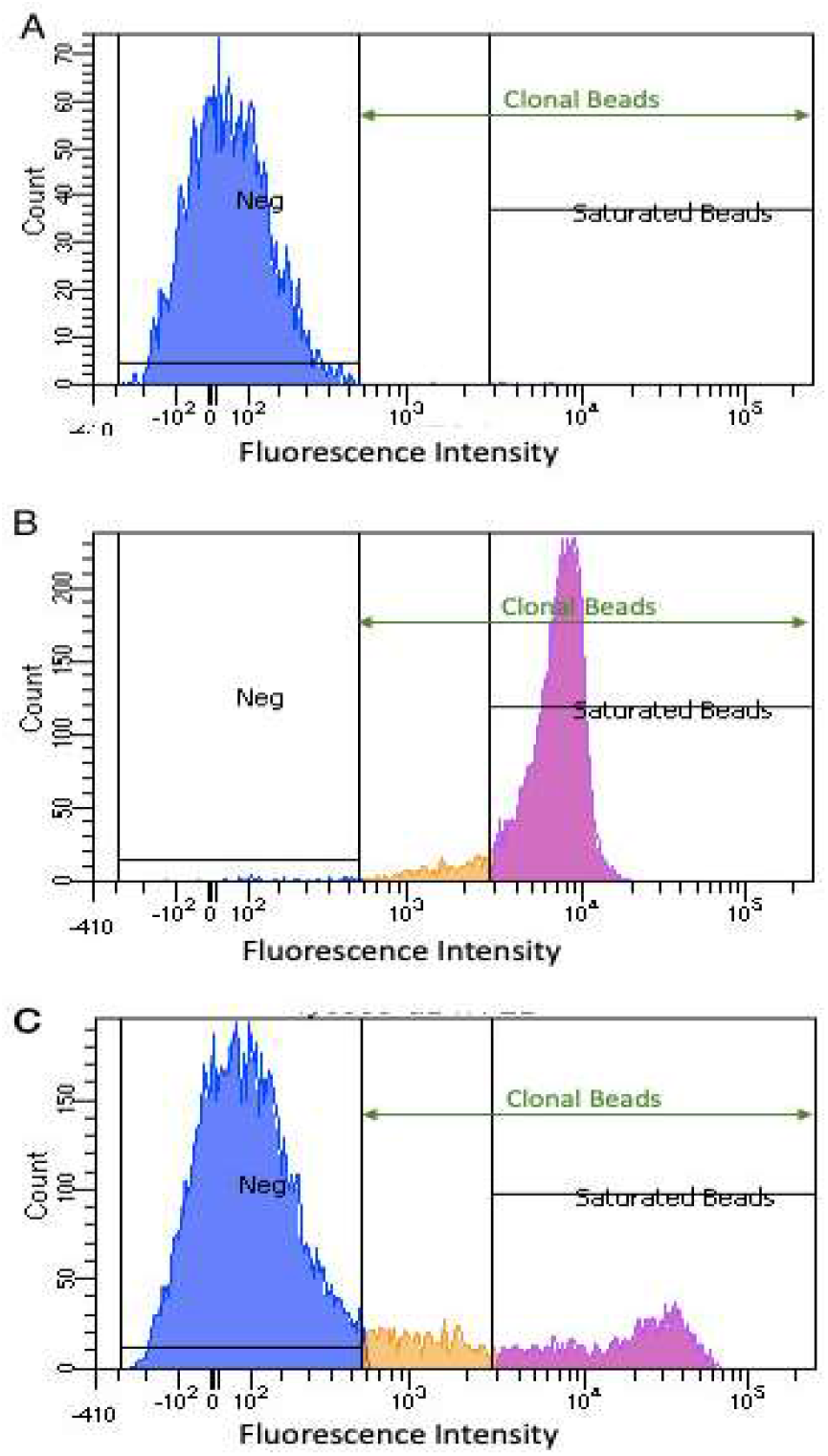
Flow cytometry gating strategy. (A) Blue negative control population defined by signal from bPCR performed without template (B) Purple positive control population defined by signal from top 90% of samples from optimised bPCR protocol(C) Example data from an on-bead ePCR sample. The proportion of ePCR beads with no measurable PCR products is gated blue. Clonal beads are indicated by fluorescence signals higher than those for the negative population. The purple “saturated” population have signals equal or greater than that of the positive control population in B.

**Figure 2.**
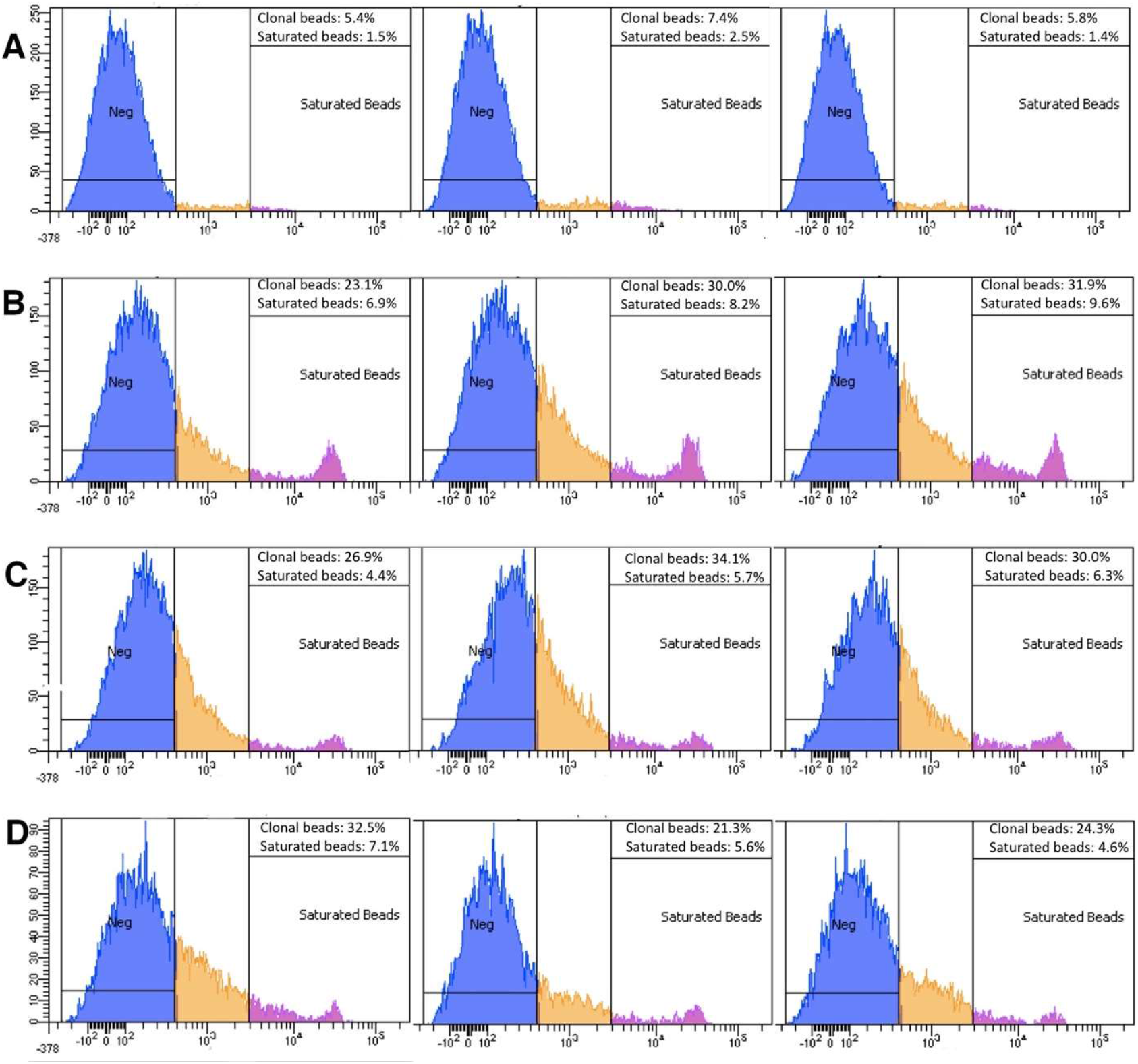
Flow cytometry histograms showing the fluorescence signals from ePCR products performed with the combinations of reagent concentrations specified in Table 1. Data from a replicate experiment are shown in the supporting information.

### 3.3 Increasing polymerase concentration improves ePCR yield

The concentration of polymerase is the first biochemical parameter we tested for emulsion PCR. Studies have revealed that adsorption of polymerase at the oil-water interface can hinder efficient ePCR.[27–29] This problem is particularly prominent when the concentration of polymerase is low; protein molecules adsorbing to the interface at low areal density can make multiple contacts with the interface, further denaturing and deactivating them.[30,31] We used the low polymerase concentration that is generally sufficient for regular PCR suggested by the supplier (i.e. 0.025 U/μl) as the reference point, and increased the concentration to 0.5 U/μl, a concentration used in some published methods for ePCR,[31,15] and assessed the change in yield and population of clonal beads. Our results (**Figure 3A and 3B)** show that increasing the concentration of polymerase increased the ePCR yield 4-fold - from 16.6 ± 0.1 to 18.7 ± 0.4, and the fraction of clonal and saturated beads increased by 15% - from 5.1%± 0.6% to 20.1%± 4.0%, and 6% - from 1.4% to 0.3% respectively, as shown in **Figure 4A and 4B**. There are several possible reasons for this phenomenon. Apart from the aforementioned denaturation, polymerase molecules will be divided amongst drops with a Poisson distribution, meaning that polymerase concentration varies between drops. Thus, excess polymerase is needed to ensure a sufficient number of enzyme molecules in all template-containing drops. Due to this variation in the number of polymerse molecules per-drop, template-containing drops with abundant polymerase will likely result in saturated clonal beads whereas others will yield clonal beads with low to medium coverage. This variation partially accounts for fewer saturated beads from ePCR than from non-emulsion bPCR control. Drops with insufficient amounts of any PCR reagent will suffer from low amplification efficiency.

**Figure 3.**
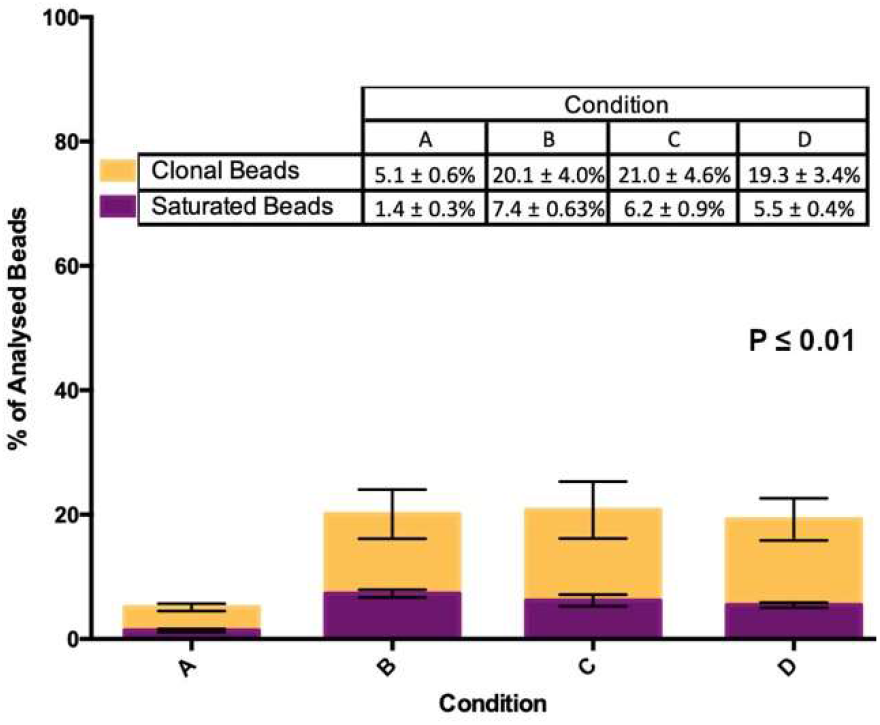
Percentage clonality and saturation of ePCR beads as determined by flow cytometry. P-value obtained from ANOVA.

**Figure 4.**
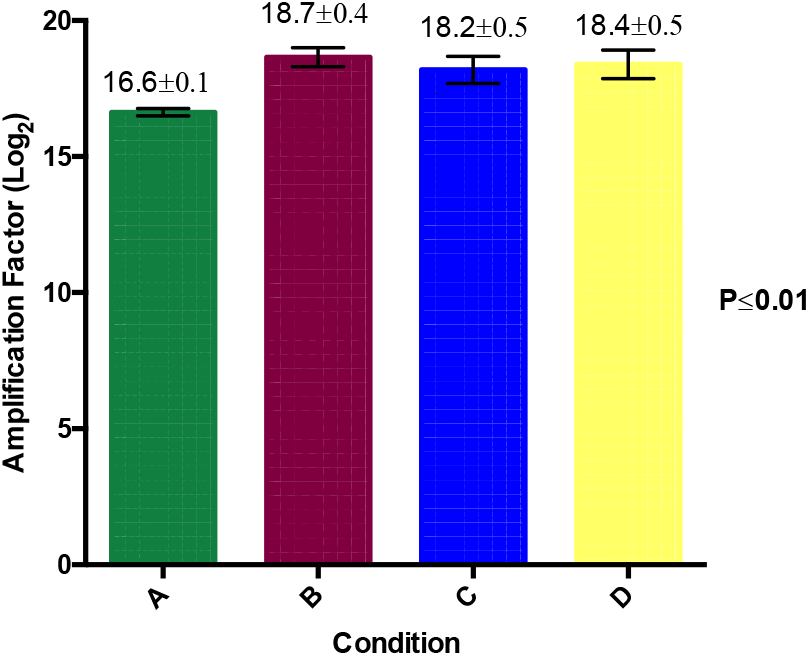
ePCR yield calculated as amplification factor in log2 scale determined by qPCR. P-value obtained from ANOVA.

### 3.4 Increasing reverse primer and dNTP concentration has no effect on ePCR yield

The results of the previous section show that there is a population of beads with amplicons on their surface but fall short of saturation. Usually, the end point of amplicon generation is attributed to the depletion of reagents like primers and nucleotides. Therefore, we generated ePCR samples with higher reverse primer concentration (10 μM vs 2 μM) and higher dNTP concentration (3.5 mM vs 0.4mM).

The data show that increasing the concentration of reverse primer has no significant effect on the percentage of clonal and saturated beads produced by ePCR (**Figure 3A and 3C**). The fraction of clonal beads produced using 2 μM reverse primer was 20.1% ± 4.0% and 20.7% ± 4.6% for 10 μM reverse primer. Similarly, the fraction of saturated beads produced using 2 μM and 10 μM reverse primer was 7.4% ± 0.6% and 6.2% ± 0.9% respectively. The amplification factor determined by qPCR was also similar for the two conditions; 18.7 ± 0.4 for the 2μM reverse primer ePCR and 18.2 ± 0.5 for the 10μM reverse primer ePCR (**Figure 4A and 4C**).

As shown in **Figure 3A and 3D**, increasing the concentration of nucleotides from 0.4 mM to 3.5 mM did not significantly increase the percentage of clonal or saturated beads. The reactions with 0.4 mM dNTP emulsion gave 20.1%± 4.0% clonal beads, similar to the 19.3% ± 3.4% clonal beads generated by the 3.5 mM dNTP emulsion. The percentage of saturated beads was similar across the two conditions as well; 7.4% ± 0.6% for 0.4mM dNTP and 5.5% ± 0.40% for 3.5 mM dNTP. The amplification factor as determined by qPCR was also similar for the two conditions; 18.7 ± 0.4 for the 0.4mM dNTP ePCR and 18.4 ± 0.5 for the 3.5mM dNTP ePCR (**Figure 4A and 4D**).

To summarize, we have examined the effects of adjusting biochemical components in ePCR on the amplification yield, and relative populations of clonal and saturated beads. The outcomes of these comparisons are summarized in Table 2. We demonstrated that a higher concentration of polymerase, 0.5 U/μl, is necessary for ~20% of the final bead population to be clonal, whereas higher concentrations of reverse primer and nucleotides did not contribute to significant improvement.

**Table 2.**
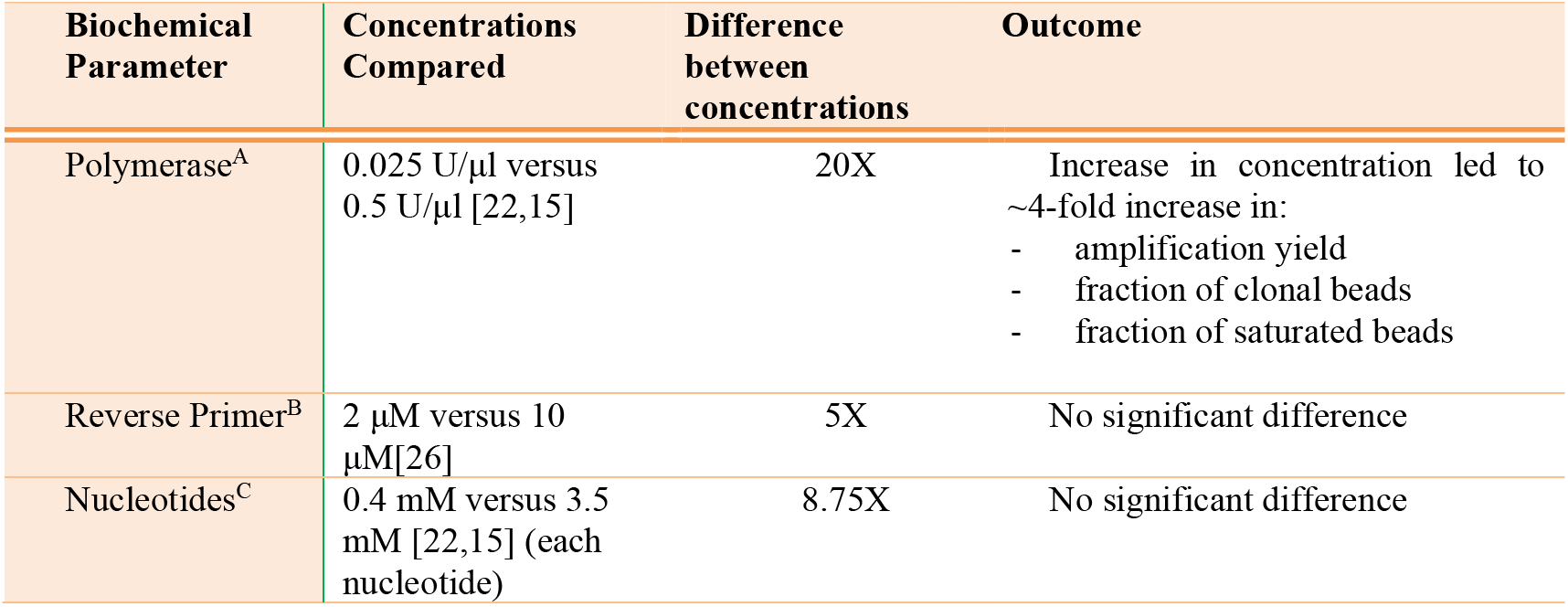
Summary of all the biochemical parameters tested and their outcomes. (A) The effect of increasing the polymerase concentration by 20-fold was tested in emulsions with 2 μM reverse primer and 0.4 mM of each dNTP. (B) The effect of increasing the reverse primer concentration 5 times was tested in emulsions with 0.5 U/μl polymerase and 0.4 mM of each dNTP. (C) The effect of increasing the nucleotide concentration 8.75 times was tested in emulsions with 0.5 U/μl polymerase and 2 μM reverse primer.

### 3.5 Discussion

Our results indicate that reverse primer and dNTP are not the limiting reagents in ePCR and adjustments elsewhere may be more effective for increasing the proportion of saturated beads. It may be worth investigating if these concentrations could be lowered without affecting amplification yield and percent clonality to avoid non-specific amplification due to excess reagents. Other factors worth investigating to further optimize ePCR performance could be the uneven distribution of beads in drops, unequal drop sizes, and even polymerase inhibition in later PCR cycles. Disentangling the contributions from each of these factors present interesting opportunities for future investigation and are worth considering to increase the user-friendliness of ePCR for different applications.

Our strategy expanded upon existing workflows to generate ‘monoclonal’ beads via on-bead ePCR with libraries of DNA templates.[6,14,16,22,32] On-bead ePCR has enabled high-throughput screening for interactions between transcription factors and DNA[16], aptamer selections,[15,22,33] detection and quantification of rare allelic mutations,[32] and high-throughput next-generation sequencing (NGS),[6,34–36] among myriad applications. While these works successfully generated monoclonal beads and demonstrated downstream utility, none of them used a single particle strategy such as ours to assess ePCR in terms of the percentage of saturated beads and amplicon coverage per bead. We have thus provided a robust approach to evaluate ePCR products. Specifically, we evaluated the extent of amplification on each ‘monoclonal’ bead. This is key for accurate and unbiased detection of sequences or sequence-dependent functions in these applications by ensuring the difference in signal between beads is sequence-specific and not due to the difference in sequence abundance. This ePCR recipe can be applied immediately by any lab and the reaction volume is scalable to accommodate the complexity of the input DNA library. This formulation can also be used without microbeads when immobilizing the library is undesired. The current study surveys the reagent concentrations recommended in the literature and finds the optimum; this has not been systematically documented previously.

## 4. Conclusion

We have contributed a single-particle strategy and systematic study to measure the influence of different concentrations of PCR components and to identify those conditions most likely to produce complete, unbiased, and non-chimeric amplicons. In this strategy, on-bead amplicons are derived only from the co-occurrence of template molecules, beads, and PCR reactants within the same drop. The distribution of blank, clonal, and saturated beads therefore reflects the situation within the emulsion drops during amplification. This analysis can resolve the inherently heterogeneous distribution of DNA templates among drops and assess whether the achieved distribution matches the one predicted by Poisson statistics to maximize the yield of monoclonal beads. This information is inaccessible by ensemble or bulk characterization of end-point products, and detailed analyses on emulsions generated by user-friendly bulk methods have not been reported. Furthermore, this high-throughput flow-cytometry approach can count orders of magnitude more ePCR products than possible by imaging drops in the same amount of time, as done in digital droplet PCR, rendering more statistical significance to the results.

We employed this approach to measure the influence of tuning the concentrations of polymerase, reverse primer, and nucleotides on the distribution and relative extent of amplification in on-bead ePCR. In doing so, we revealed viable starting parameters for ePCR. The concentrations we chose to test have been published individually by other researchers but direct comparisons of the different combinations had not been reported. Experiments require choosing a concentration for each component and our data enable researchers to make evidence-based decisions. Our data revealed outcomes contrary to some published recommendations, as summarized in **Table 2** and detailed in **Figures 3 and 4**. Specifically, the 20-fold increase in polymerase to 0.5 U/μl was necessary to achieve sufficient yield, but it was unnecessary to increase the concentration of reverse primer and each nucleotide beyond 2 μM and 0.4 mM, respectively. These results and the emulsion formula and generation conditions reported here can be adapted immediately by any laboratory with these common reagents and standard laboratory equipment, such as stir plates and pipettes. The conclusions of this study are equally insightful for ePCR with or without beads; the beads enable the analysis performed herein and select applications but are not required for successful ePCR. Our results therefore increase the accessibility of ePCR to newcomers and facilitate the integration of ePCR into applications such as liquid biopsy and cancer research,[37,38] metagenomic studies,[39,40] and aptamer discovery.[5,15,18,22,26]

## Author contributions

The manuscript was written through contributions of all authors. All authors have given approval to the final version of the manuscript.

## Credit author statement

Ryan H.P. Siu: Investigation, Methodology, Writing – Original draft. Yang A. Liu: Investigation, Methodology, Formal analysis, Writing – Original draft. Kaitlin H.Y. Chan: Investigation, Methodology, Writing – Original draft, Visualization. Clara Ridzewski: Investigation, Methodology, Visualization. Liane Siu Slaughter: Conceptualization, Funding Acquisition, Supervision, Project Administration, Investigation, Methodology, Writing – Reviewing and Editing Angela Wu: Conceptualization, Funding Acquisition, Supervision, Writing –Reviewing and Editing.

## Appendix A: Supporting Information

The Supporting Information to this article can be found online at ___

The Supporting Information includes the sequences of the DNA template, primers and probes, experimental details and results of the non-emulsion bead-based PCR, predictions based on the Poisson distribution model, MgCl_2_ titration experiment and droplet size distribution of two oil formulations

## Acknowledgements

We thank Professor Shuhuai Yao, Dr. Xiaonan Xu, and Mr. Binbin Cui for aliquots of ABIL^®^ EM 90 and assistance with characterization in early phases of the project. We thank Professors Danny Leung, Tom Cheung, and Andrew Miller and their research groups, and the support staff of the Biosciences Central Research Facility at HKUST. We also thank Professor Tom Soh and Dr. Jianpeng Wang for helpful discussions. This work was funded by HKUST’s start-up and initiation grants (Hong Kong University Grants Committee), the Hong Kong RGC Early Career Support Scheme (RGC ECS 26101016), the Hong Kong Innovation & Technology Commission Innovation and Technology Fund (ITS/350/16) and the Hong Kong Epigenomics Project (LKCCFL18SC01-E). Liane Siu Slaughter is grateful for support through a Junior Fellowship from the Hong Kong Jockey Club Institute for Advanced Study, The Hong Kong University of Science and Technology (2016-2018).

