## Supplementary Information for "Optimization of On-Bead Emulsion Polymerase Chain Reaction Based on Single Particle Analysis"

*corresponding authors

This supporting information includes one table of all sequences used in this study, and eight figures showing the results from non-emulsion bead-based PCR, tests of emulsion stability of two published formulations, predicted distributions of DNA templates according to the Poisson model, and histograms from flow cytometry analysis of samples from replicate ePCR experiments of those discussed in the main text, data from non-emulsion on-bead PCR (bPCR) experiments with different $MgCl_2$ concentrations and PCR cycle numbers, and droplet size distribution analysis of microscope pictures of emulsion before and after heat cycling.

* Corresponding author

Present Addresses:
∫ *Phase Scientific International; 32&33/F, Gravity, 29 Hing Yip Street, Kwun Tong, Kowloon, Hong Kong*
‡ *Department of Molecular Genetics, University of Texas Southwestern Medical Center, Dallas, TX 75390, USA*
†*Biotechnology Centrum BIOTEC, TU Dresden, Tatzberg 47/49.01307 Dresden, Germany*
¥ *Clear Water Bay, Hong Kong*

**Table S1. Sequences of oligonucleotides used in this study**

| | |
|---|---|
| Template | |
| H4K16ac aptamer template | 5′-AGCCTCAAGATCATCAGCAAAGACGTAAGTTAATTGGACTTGGTCGTGTGCGGCACAGCGATTGAAATTTGGTATCGTGGAAGGACTC-3’ |
| Primers | |
| forward primer (GD1F) | 5′-AGCCTCAAGATCATCAGCAA-3′ |
| reverse primer (GD1R) | 5′-GAGTCCTTCCACGATACCAA-3′ |
| Amino modified GD1F | 5’-amino-PEG$_{18}$- AGCCTCAAGATCATCAGCAA-3′ |
| Probes | |
| Tye665-GD1F reverse complement | 5' TYE™ 665 – TTGCTGATGATCTTGAGGCT-3′ |
| Tye665-GD1R | 5' TYE™ 665 – GAGTCCTTCCACGATACCAA-3′ |

**Non-emulsion bead-based PCR (bPCR) protocol and result**

Each bPCR consisted of 1X GoTaq PCR Master Mix, 25mM $MgCl_2$, 0.025U/µl G2 Hot-Start polymerase, $3x10^5$ FP-beads, 2 µM 5' TYE™665–GD1R and 10 nM template. We then performed bPCR under the following cycling conditions: 95 °C for 3 min, followed by varying cycles of 95 °C for 20 sec, 65 °C for 30 sec and 72 °C for 1 min and a post-cycling extension at 72 °C for 5 min.

Multiple PCR blocks were used to ensure that PCR samples undergo their designated number of cycles and the 5 min post-cycling extension. To promote efficient amplification, samples were vortexed briefly every 3 to 4 PCR cycles to avoid beads aggregation.

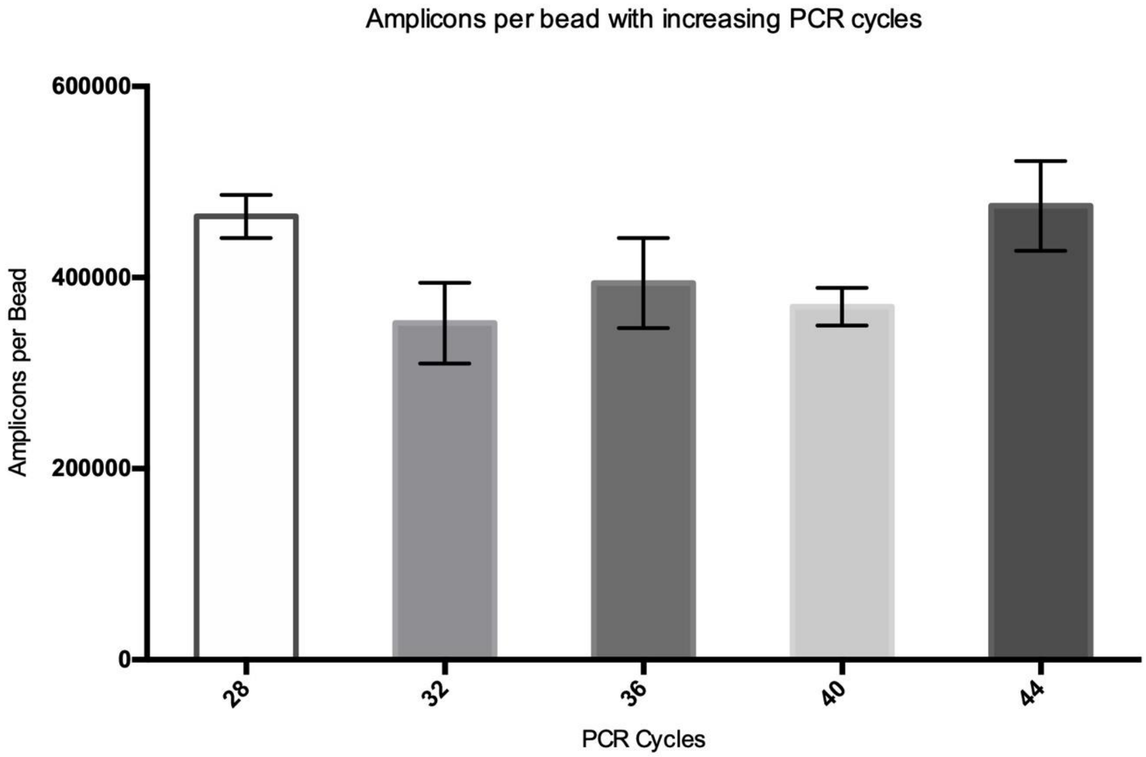


**Figure S1.** Amplicons per bead from bPCR samples subjected to different numbers of PCR cycles. The GD1 sequence and primers were used for this test. Amplicon per bead is calculated from the number of amplicons quantified by qPCR divided by number of beads subjected to the assay.

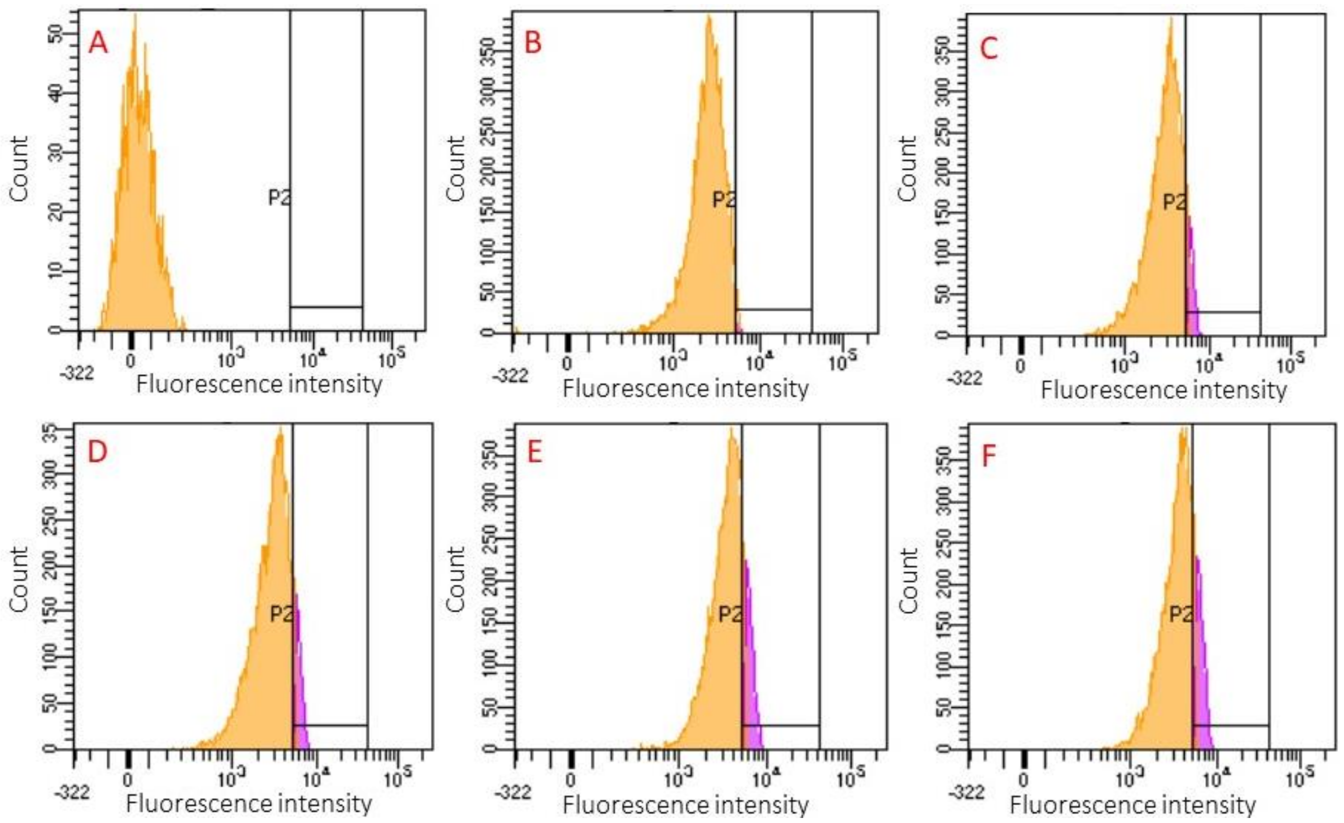


**Figure S2.** Histograms of the fluorescence signal from singlet beads obtained by flow cytometry. Data are from (A) unlabeled FP-beads and labeled beads from bPCR with (B) 28 cycles, (C) 32 cycles, (D) 36 cycles, (E) 40 cycles, (F) and 44 cycles of bPCR products. The fluorescence intensity is directly proportional to the number of amplicons per bead. The beads underwent the same dehybridization, hybridization, and characterization workflow as beads isolated from ePCR. The median fluorescence intensity of the six plots are 8, 2222, 2767, 2758, 3252, 3287 arbitrary units (au) respectively. The populations shaded in purple represent beads where the number of amplicons approaches the maximum number limited by the forward primers on the bead. This criterion is established from the fluorescence of FP-beads hybridized with dye-labeled forward primer reverse complement.

When increasing cycle number for bPCR from 28 to 44, the median fluorescence intensity in Figure S2 increases from 2222 to 3287 au. However, qPCR derived data in Figure S1 does not indicate such that the number of amplicons per bead changes. Since the data shows that 28 cycles are enough to produce a narrowly distributed population of beads with sufficient number of amplicons per bead during bPCR, we chose to use this cycle number to produce reference positive control beads. These data from non-emulsion bead-based PCR serve as a reference to which to compare the data from ePCR tests.

**Emulsion stability of two oil formulations (EM90-based and Span-80-based)**

We chose to compare two popular oil phase formulations comprising easily obtainable off-the-shelf reagents, which we call ‘EM90-based’ and ‘Span80-based’. These formulations are widely published, but there are few side-by-side comparisons of how the emulsions fare over the complete course of PCR heating programs.[18,19,36–38] We generated emulsions with PCR buffer and FP-beads in the aqueous phase with two different oil formulations. After subjecting the emulsions to the PCR cycling program described in the main text for 10 cycles, we recorded the macroscopic appearance of the emulsions.

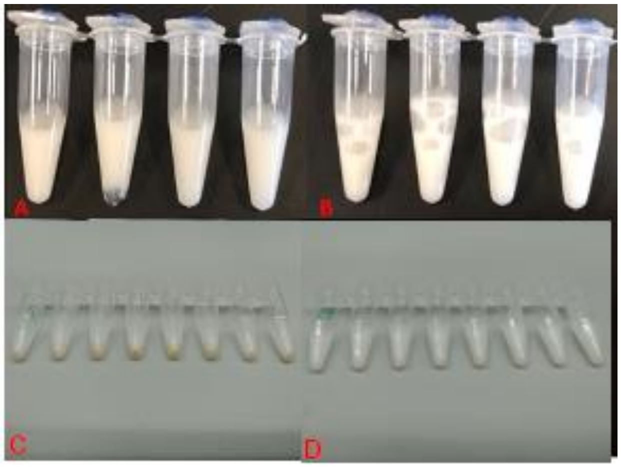


**Figure S3** Appearance of emulsions in PCR tubes. Emulsions were generated with aqueous phases containing PCR buffer and FP-beads and two different oil formulas. (A) Span80-based emulsion before PCR cycling. (B) EM90-based emulsion before PCR cycling. (C) Bottom view of Span80-based emulsion after PCR cycling. (D) Bottom view of EM90-based emulsion after PCR cycling. All photos herein were taken by the authors.

The results shown in Figure S3 served as a preliminary qualitative assessment of the thermal stability of two tested oil formulas through multiple PCR cycles. While Panels A and B appear similar in color and consistency before heating, Panels C and D show contrast in their appearance after heating. Specifically, brown dots are observed at the bottoms of the tubes for Span80-based emulsions shown in Panel C, but not for the EM90-based emulsions in Panel D. These brown dots are aggregates of beads that likely result from coalescence of emulsion drops. To further verify the thermal stability of the two formulas, microscopic images were taken in Figure S4 to confirm any bead aggregation.

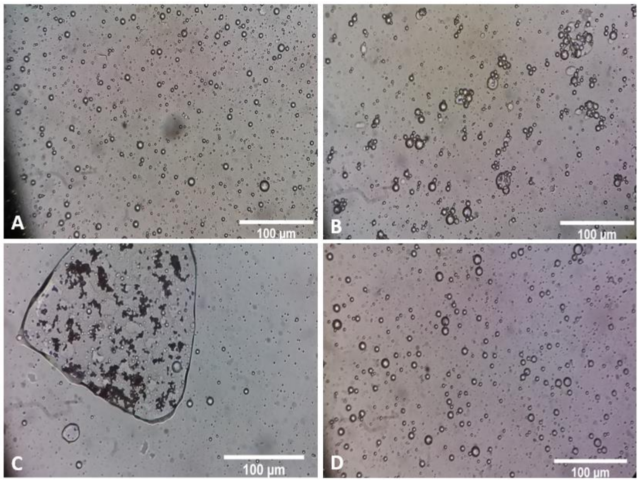


**Figure S4** Representative microscope images (40X magnification) of drops before and after heating the emulsions with the two oil formulas. Samples were taken from the bottom of the emulsions and diluted in 10 times the volume of mineral oil before transferring to a coverslip. (A) Span80 emulsion prior to heating. (B) EM90-based emulsion prior to heating. (C) Span80 emulsion after heat cycling shows merged aqueous droplets in oil with large clusters of magnetic beads. (D) EM90-based emulsion did not undergo visible morphological changes after heat cycling.

**Figure S4** shows microscopic images of the emulsion samples taken from conditions stated in **Figure S3**. As shown in Panels B and D,  the EM90-based droplets had no morphological difference throughout PCR cycling. However, a cluster of beads encapsultated in a large drop can be observed in the Span80-based emulsion after heating as shown in **Panel C.** This is not in visble in the emulsion before heating (**Panel A)**. This observation indicates that the drops in the Span80-based emulsion had fused together during heating. We therefore ruled out using the Span80-based oil formula and use only the EM90-based formula for the remainder of this study.

**Poisson distribution model**

The distribution of DNA molecules into separate droplets generally follows the Poisson distribution. We adopted 2 pM of DNA template as the starting input concentration, as it is the concentration used for the studies herein and prior literature, and used the Poisson model to predict the proportion of droplets carrying a specified number, k, of template molecules.

$$P(k, \lambda) = e^{-\lambda}\frac{\lambda^k}{k!}$$

k = number of DNA template per drop

λ = Average number of template per drop

P(k, λ) = Probability of having k DNA template per drop

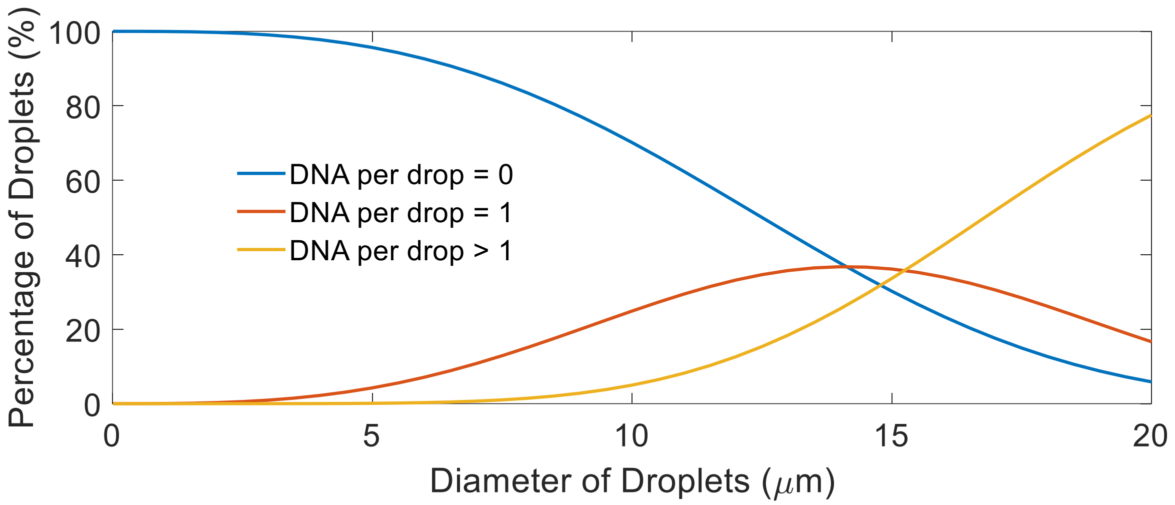


**Figure S5.** Percentage of droplets carrying zero, one, or more than one DNA template per drop for different average droplet diameters and 2 pM of input DNA. In the ideal distribution for ePCR, the percent of drops having one template molecule (k = 1) is maximized, to maximize the number of productive reaction compartments. Simultaneously, the percent of drops having multiple template molecules (k > 1) is minimized, to minimize the occurrence of chimeric products. Based on this model, droplets with diameters of 7 to 10 μm adequately satisfy these criteria. We therefore tuned the stirring speeds and times to target average drop sizes within this window of diameters.

### On-bead ePCR Histograms from Flow Cytometry

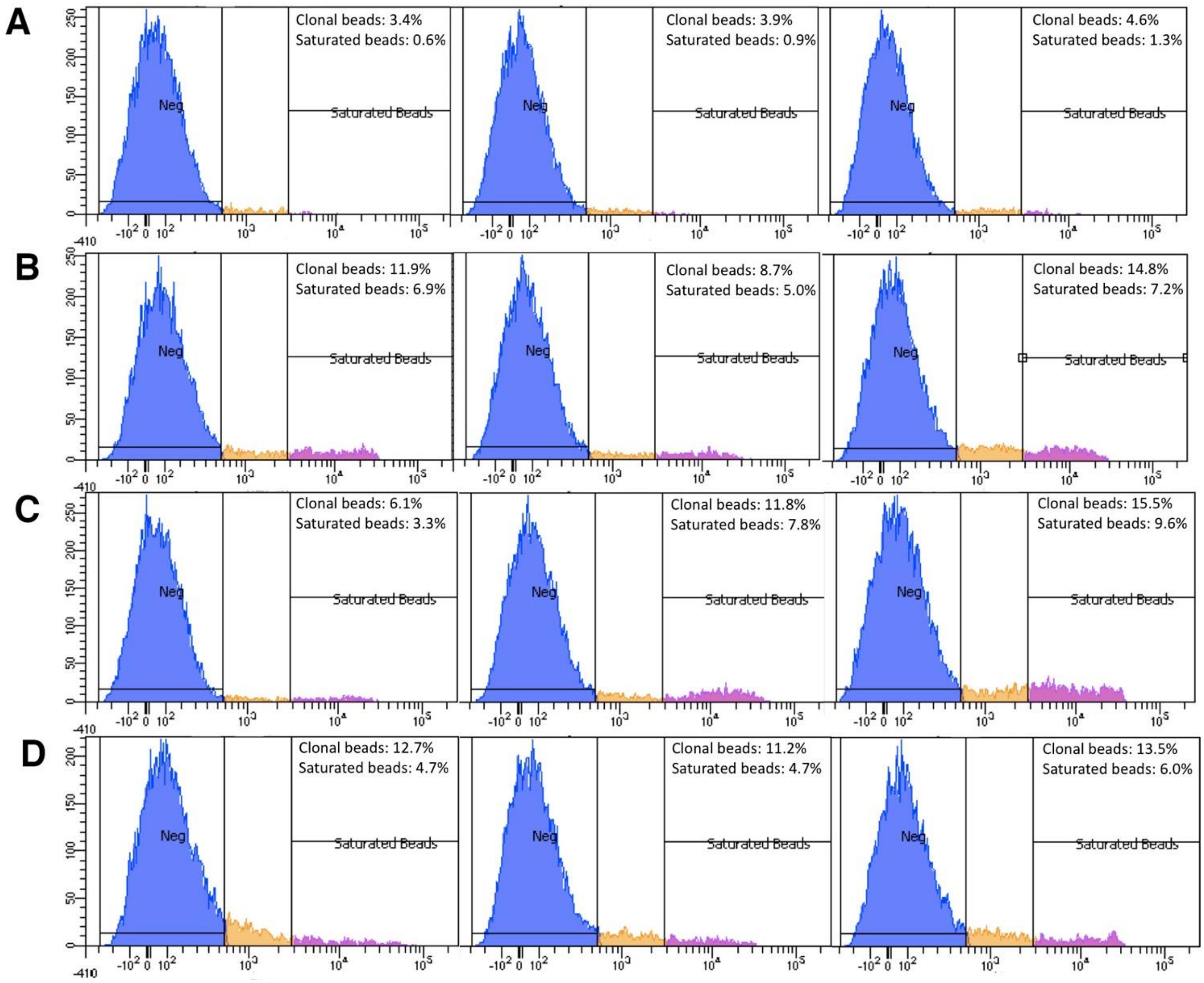


**Figure S6.** Histograms of ePCR products performed with different combinations of reagent concentrations, a replication of the experiment for the data shown in Figure 2. All ePCR were performed with the 'EM90-based' oil formula. The combinations follow those tabulated in Table 1. (A) 100 ul of PCR aqueous phase with 2 μM reverse primer, 0.4 mM dNTP each and 0.025 U/μl polymerase. (B) 2 μM reverse primer, 0.4 mM dNTP each and 0.5 U/μl polymerase. (C) 10 μM reverse primer, 0.4 mM dNTP each and 0.5 U/μl polymerase. (D) 2 μM reverse primer, 3.5 mM dNTP each and 0.5 U/μl polymerase.

**Testing the influence of different $MgCl_2$ concentrations with non-emulsion bead-based PCR**

We performed an experiment to assess the effect of $MgCl_2$ concentration on PCR yield. We used the the non-emulsion bead-based PCR described in the Methods with 2mM $MgCl_2$ and 25mM $MgCl_2$.

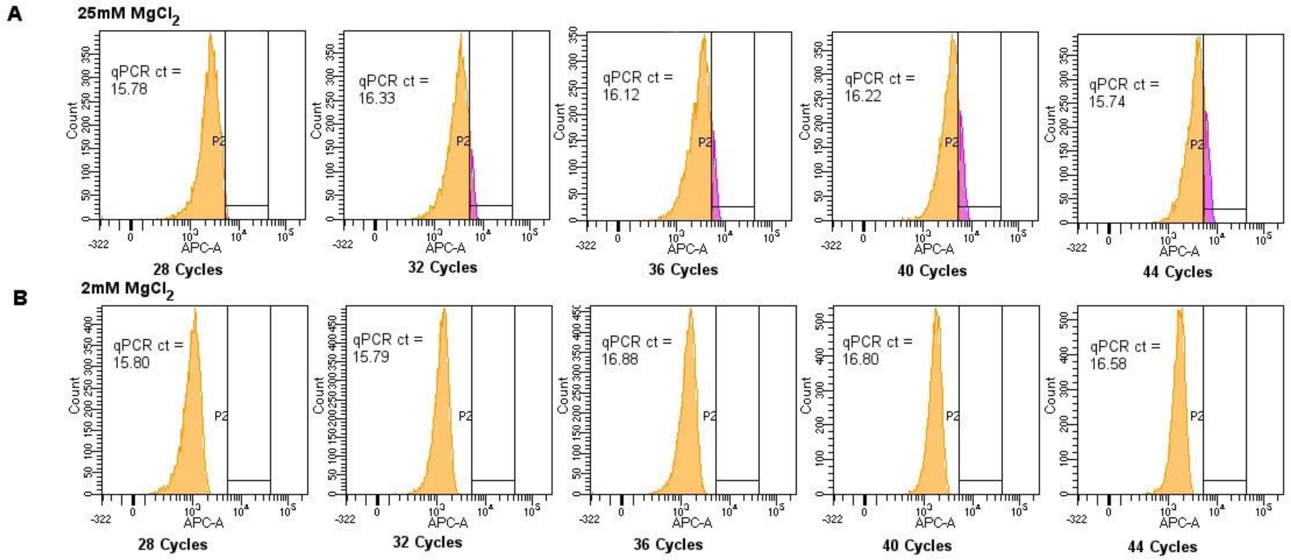


**Figure S7.** Histogram of bPCR products produced with 2mM or 25mM $MgCl_2$ bPCR settings with increasing PCR cycles.

The flow cytometry results show that 25mM $MgCl_2$ bPCR consistently produces more beads with more amplicons than bPCR with 2 mM $MgCl_2$. To further verify if the products are the intended amplicon, we checked the melting curves of the on-bead products.

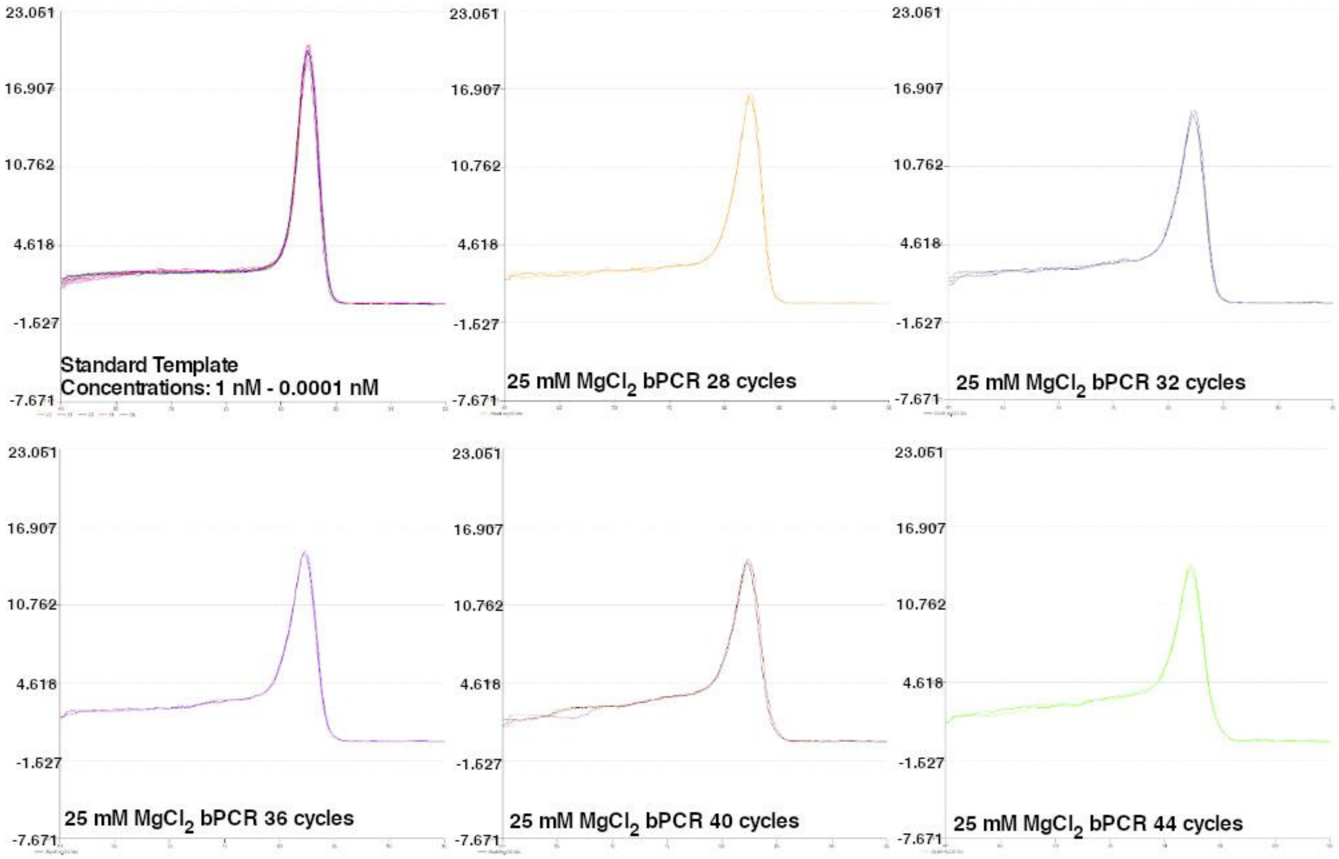


**Figure S8.** Melting curves of standard template and bPCR products. The melting temperature for the standards range from 82.39 °C to 82.59°C and the melting temperature for the bPCR products range from 81.95°C to 82.39°C.

### Droplet size distribution

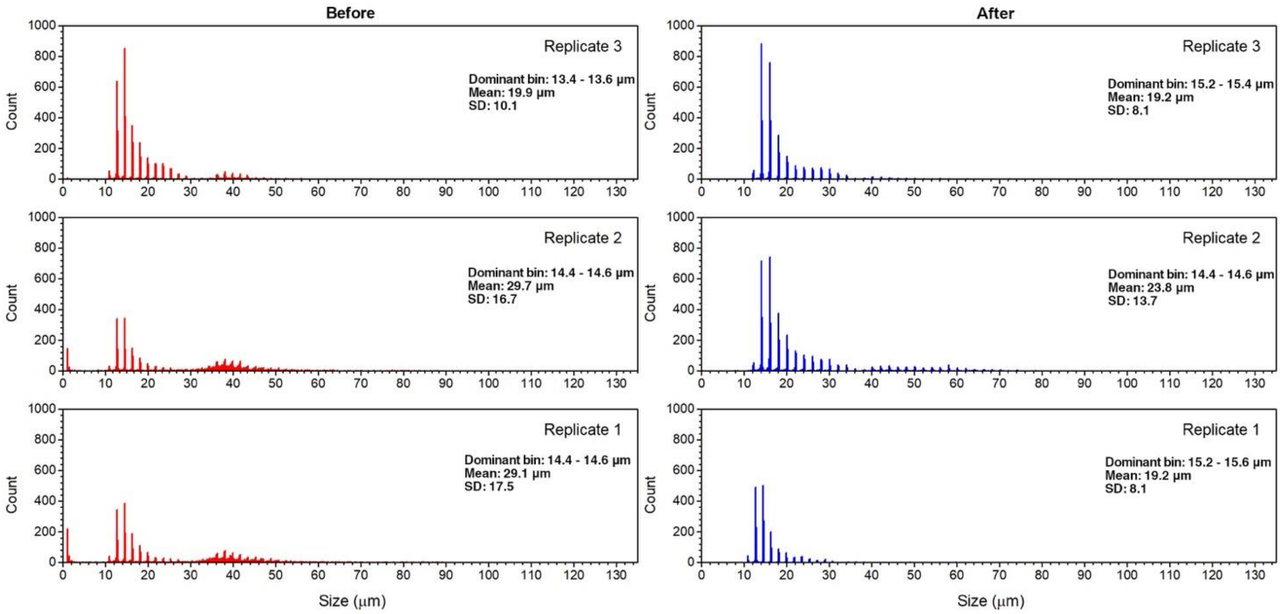


F**igure S9.** Histogram of size distribution of emulsion droplets before and after 50 PCR cycles. `

We quantitatively characterized the size distribution of droplets in the emulsion using MIPAR. We found that despite the wide range of sizes, a large majority of the droplets were close to 14.5 μm in diameter. Furthermore, the size distribution profiles of the emulsions before and after heat cycling are very similar. This indicates that the emulsion was indeed stable throughout ePCR and if the stirring conditions are kept constant, emulsion reproducibility is possible.
